# CELF2 promotes tau exon 10 inclusion via hinge domain-mediated nuclear condensation

**DOI:** 10.1101/2024.11.02.621395

**Authors:** Xin Li, Ishana Syed, Zhao Zhang, Rashmi Adhikari, Dan Tang, SuHyuk Ko, Zhijie Liu, Lizhen Chen

## Abstract

Alternative splicing is a fundamental process that contributes to the functional diversity and complexity of proteins. The regulation of each alternative splicing event involves the coordinated action of multiple RNA-binding proteins, creating a diverse array of alternatively spliced products. Dysregulation of alternative splicing is associated with various diseases, including neurodegeneration. Here we demonstrate that CELF2, a splicing regulator and a GWAS-identified risk factor for Alzheimer’s disease, binds to mRNAs associated with neurodegenerative diseases, with a specific interaction observed in the intron adjacent to exon 10 on Tau mRNA. Loss of CELF2 in the mouse brain results in a decreased inclusion of Tau exon 10, leading to a reduced 4R:3R ratio. Further exploration shows that the hinge domain of CELF2 possesses an intrinsically disordered region (IDR), which mediates CELF2 condensation and function. The functionality of IDR in regulating CELF2 function is underscored by its substitutability with IDRs from FUS and TAF15. Using TurboID we identified proteins that interact with CELF2 through its IDR. We revealed that CELF2 co-condensate with NOVA2 and SFPQ, which coordinate with CELF2 to regulate the alternative splicing of Tau exon 10. A negatively charged residue within the IDR (D388), which is conserved among CELF proteins, is critical for CELF2 condensate formation, interactions with NOVA2 and SFPQ, and function in regulating tau exon 10 splicing. Our data allow us to propose that CELF2 regulates Tau alternative splicing by forming condensates through its IDR with other splicing factors, and that the composition of the proteins within the condensates determines the outcomes of alternative splicing events.

## INTRODUCTION

Alternative splicing (AS) plays an essential role in transcriptional regulation and is found in approximately 95% of human protein-coding genes. AS gives rise to multiple mature mRNAs encoding protein isoforms with distinct functions, structures, stability, or cellular localization across different tissues, developmental stages, or under specific conditions ^1^. The existence of distinct splice isoforms suggests unique temporal and spatial functionalities, thereby implicating AS in fundamental biological processes as well as disease pathogenesis ^2^. AS of pre-mRNA is carried out by the spliceosome, a macromolecular complex comprising small nuclear RNAs and small nuclear ribonucleoproteins (snRNPs). Within the regulatory framework, AS-related RNA-binding proteins (RBPs) play an instrumental role in regulating AS events. Sequence-specific RBPs binds to pre-mRNA to form ribonucleoprotein complexes to control AS, and each AS event is controlled by multiple RBPs.

To achieve precise spatiotemporal control over intricate biochemical reactions, cells must orchestrate the arrangement of proteins and other large molecules within subcellular regions. Beyond traditional membrane-bound organelles like the endoplasmic reticulum and Golgi apparatus, cells harbor diverse membraneless compartments. These include nucleoli, Cajal bodies, processing bodies and stress granules ^3–5^. Recent research suggests that the formation of these membraneless compartments is often mediated by a physical phenomenon called phase separation (PS). And PS has emerged as a mechanism underlying spatiotemporal protein recruitment involved in various physiological processes, including splicing regulation, transcription, signal transduction, and DNA damage repair ^6–10^. Among PS proteins, many have intrinsically disordered regions (IDRs) that lack a specific three-dimensional structure and typically are enriched with charged amino acids, polar amino acids, and/or aromatic amino acids. Multivalent electrostatic, cation-pi, pi-pi, and hydrophobic interactions have all been proposed to contribute to IDR PS ^11–17^.

RBPs are generally characterized by the RNA-recognition motifs (RRMs) and IDRs, which allow RBPs to assemble with RNAs and proteins to form dynamic phase-separated condensates. RBPs are critical players in gene regulation and are at center stage in our understanding of cellular function in both normal and disease processes. Dysfunction of RBPs and the subsequent disruption of RNA processing are increasingly implicated in neurological disorders including age-associated neurodegeneration ^18^. CELF genes encode a conserved family of RBPs that is involved in regulating gene expression at multiple levels, including alternative splicing, RNA transport, translation and mRNA stability ^19^. All 6 human CELF genes are expressed in both developing and adult brains, and the expression of CELF genes in the nervous system is evolutionarily conserved ^20–25^. CELF2, like the other five members of its family, possesses three RNA recognition motifs (RRMs) and intrinsically disordered regions (IDRs), and is known to regulate alternative splicing ^26,27^. SNPs of the CELF2 gene have been found to be associated with late onset Alzheimer’s disease (AD) ^28,29^, offering valuable first insight into the genetic connection. However, the mechanistic role that CELF2 may play in AD is largely unknown.

A prominent pathological feature in AD and multiple neurodegenerative diseases is the aberrant aggregation of the microtubule associated protein tau (MAPT). Through AS, MAPT gene generates six protein isoforms that differ by the presence or absence of two N-terminal domains (N1 and N2) and the second microtubule binding repeat (R2) ^30^. The inclusion or exclusion of exon 10 that encodes R2 produces 4-repeat (4R) and 3-repeat (3R) tau respectively. The major function of tau is to promote microtubule assembly and stabilize microtubules by binding to microtubules ^31,32^. The 4R tau can promote microtubule polymerization faster than 3R tau ^30^. In the healthy adult human brain, the 4R and 3R isoforms are found approximately 1:1 ratio ^30,33^. Alterations in the 4R:3R tau isoform ratio, often caused by mutations within exon 10 or mutations affecting exon 10 inclusion, lead to pathogenesis of various tauopathies ^34^. The significance of preserving the equilibrium between 3R and 4R isoforms in healthy neurons is underscored by the link between perturbations in this ratio and disease.

In this study, we demonstrate that CELF2 binds to MAPT mRNAs and promotes exon 10 inclusion. Loss of CELF2 in the mouse brain results in a reduced 4R:3R tau ratio. The intrinsically disordered hinge domain of CELF2 mediates CELF2 phase-separated condensation and function but can be substituted by IDRs from FUS and TAF15. A conserved negatively charged residue within the IDR is critical for CELF2 condensation and function. Through TurboID, we identified CELF2-interacting proteins including NOVA2 and SFPQ, which coordinate with CELF2 to regulate the alternative splicing of tau exon 10. Together, these results suggest that CELF2 regulates tau alternative splicing by forming condensates through its IDR with other splicing factors, and that the composition of the proteins within the condensates determines the outcomes of alternative splicing events.

## RESULTS

### CLIP-seq identified MAPT as a target of CELF2 and Celf2 knockout alters 4R:3R tau ratio in mouse brain

To investigate CELF2 function, we have previously performed CLIP-seq in N2A cells to identify CELF2 targets ^35^. Different cell fractions (whole cell, cytoplasmic and nuclear) were used in CLIP-seq to distinguish CELF2 function in the nucleus and cytoplasm. The resulting 4279 nuclear target genes underwent KEGG pathway enrichment analysis, highlighting the enrichment of 108 genes linked to Alzheimer’s disease (AD), including tau (**Fig. 1A, 1B**). CELF2 specifically binds to the intron 5’ to the alternatively spiced exon 10 (**Fig. 1B**). This finding aligns with previous GWAS studies that have connected CELF2 with AD ^28,29^. We found that cytoplasmic CELF2 binds to the 3’UTR of mRNA, while nuclear CELF2 binds to introns, consistent with its nuclear function in alternative splicing (**Fig. S1A**). To confirm the regulation of Tau splicing by CELF2 in brain, we examined Tau exon 10 splicing in embryonic brain of Celf2 knock out (KO) mice (**Fig. S1B**) ^35^. As previously reported ^36,37^, embryonic mouse brain expresses a low level of 4R tau, with 3R tau as the predominant tau isoform (**Fig. 1C, 1D**). Notably, the 4R tau was undetectable in *Celf2* KO embryonic brain (**Fig. 1C, 1D**). Therefore, these data indicate that CELF2 binds to the intron 5’ to exon 10 and promote axon 10 inclusion (**Fig. 1E**).

**Fig. 1.**
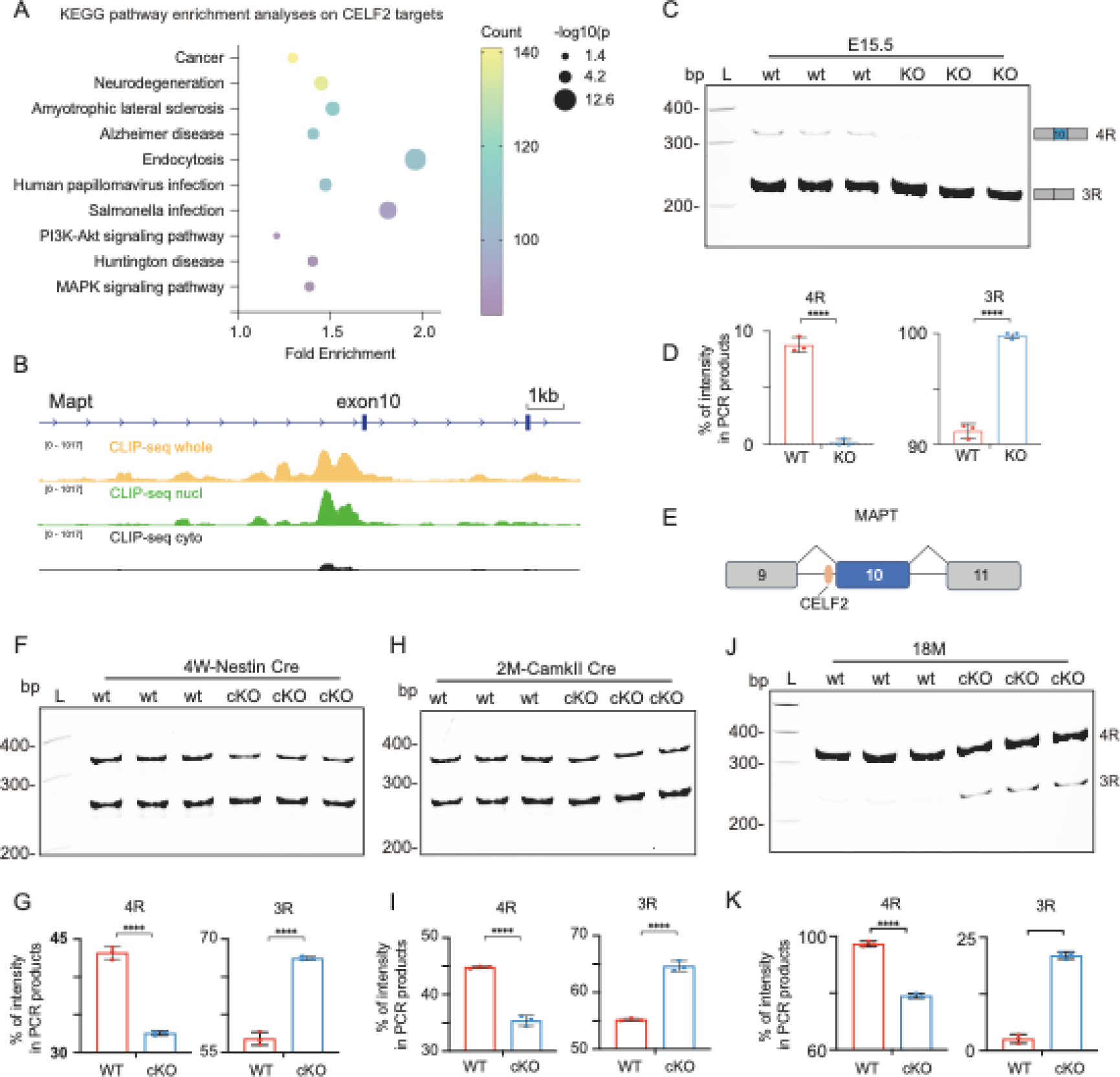
CLIP-seq identified MAPT as a target of CELF2 and Celf2 knockout reduces 4R:3R tau ratio in mouse brain. **(A)** KEGG pathway enrichment analysis on CELF2 target genes. CELF2 targets were identified from our previous CLIP-seq using N2A cells. The top 10 pathways are presented. **(B)** Genome browser view of CLIP-seq peaks on the Mapt gene locus. CELF2 specifically binds to the intron 5’-to the alternatively spliced exon 10. **(C)** RT-PCR showing that Celf2 KO eliminates 4R tau expression. Brain tissues from Celf2 KO and wildtype E15.5 mouse embryos were used to generate cDNA for RT-PCR to detect the alternative splicing of *Mapt* exon 10. **(D)** Quantification of RT-PCR results to show that Celf2 KO leads to a loss of 4R tau expression in E15.5 mouse brain. n=3, *p < 0.05; ***p < 0.001; ****p < 0.0001. **(E)** A schematic model to indicate the binding of CELF2 on the intron 5’-to exon 10 to promote exon 10 inclusion. **(F-K)** RT-PCR and quantification of 4R and 3R tau expression in mouse brains at indicated stages (4 weeks, 2 months and 18 months) in Celf2 cKO induced by Nestin-Cre (F-G) or CamKII-Cre (H-K). n=3, *p < 0.05; ***p < 0.001; ****p < 0.0001.

4R tau expression is limited in both human and rodent fetal brains ^36,37^. As postnatal development progresses, the ratio of 3R to 4R tau isoforms in the healthy human brain transitions to a 1:1 ratio ^38^. The expression patterns of tau isoforms in adult brains vary between humans and rodents. In the adult rodent brain, the presence of 3R tau gradually reduces, with 4R tau emerging as the predominant isoform in later stages of adulthood ^36,37^. To understand whether CELF2 regulates tau exon 10 splicing in adult brain, we crossed our previously generated Celf2 cKO allele with nestin-Cre or CamKII-Cre line and induced *Celf2* depletion at postnatal stages, as constitutive *Celf2* KO animals die at newborn ^35^. We confirmed the depletion of CELF2 at the mRNA and protein levels in the *Celf2* cKO brain (**Fig. S1B-H**) and examined tau exon 10 expression at 4 weeks, 2 months and 18 months old. Unlike the embryonic brain, the young adult wildtype control brains (4weeks and 2 months old) expressed approximately equal amount of 4R and 3R tau isoforms (**Fig. 1F-I**). Notably, in 18 months old brains, the 3R tau was barely detectable (**Fig. 1J, 1K**). In all stages we examined, we found that CELF2 depletion significantly reduced 4R and increased 3R tau expression (**Fig. 1F-K**), indicating that CELF2 functions to promote tau exon 10 inclusion in the brain across developmental stages.

### CELF2 forms phase-separated condensates through its intrinsically disordered hinge domain

Formation of biomolecular condensates through phase separation (PS) has emerged as a ubiquitous mechanism to promote compartmentalization for dynamic biological processes ^39–41^. Intrinsically disorder regions (IDRs) are often found in proteins that undergo PS to form condensates. Like other CELF family members, CELF2 contains an intrinsically disordered hinge domain (**Fig. S2A-C**). This prompted us to test if CELF2 undergoes PS through its IDR. We first used the optogenetic system, optoDroplet, which is based on *Arabidopsis thaliana* CRY2 ^42^. We expressed CELF2 fused to mCherry and the photolyase domain of the CRY2 protein in 293T cells (**Fig. 2A**). Upon blue light illumination, CRY2 undergoes self-association, leading to an increase of local concentration of the fused protein ^43^. Consistent with previous reports, CRY2-mCherry alone showed little clustering upon blue light activation. As expected for IDRs, fusing full length CELF2 or CELF2 IDR to CRY2 led to blue-light dependent clusters formation (**Fig. 2B, 2C**), suggesting that CELF2 IDR (the hinge domain) can drive phase-separated condensate formation. We next purified recombinant GFP-CELF2IDR (**Fig. S2D**) and performed *in vitro* droplet formation assay. Purified GFP-CELF2IDR fusion protein formed droplets in the presence of PEG8000 and the droplet size and number correlated with protein concentration (**Fig. 2D, 2E**), again suggesting that CELF2 IDR can drive condensates formation.

**Fig. 2.**
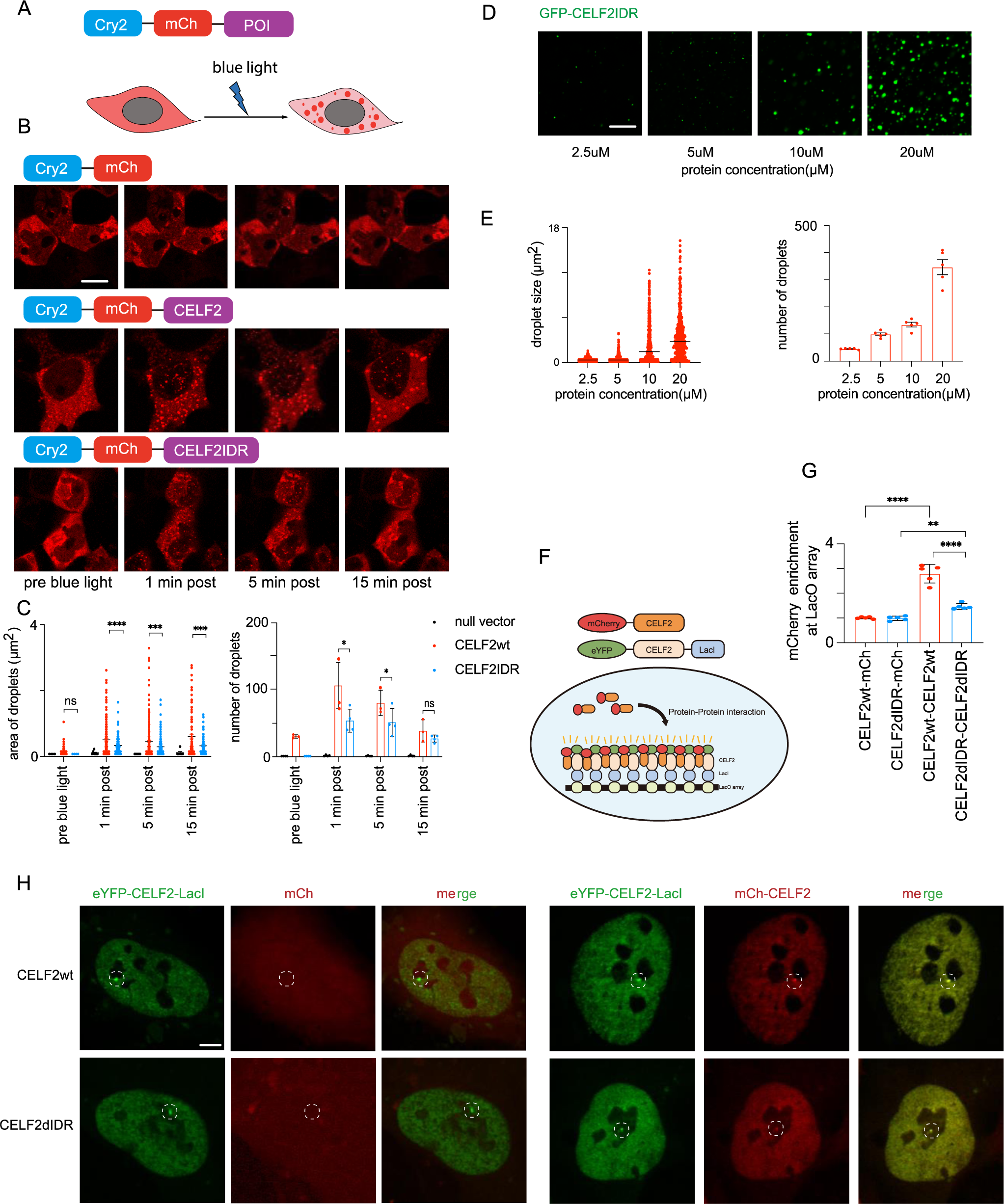
CELF2 forms phase-separated condensates through its intrinsically disordered hinge domain. **(A)** Schematic diagram of the optogenetic platform to study phase separation. The construct consists of the CRY2PHR domain, mCherry fluorescent protein and the protein of interest (POI). Upon 488nm blue light activation, CRY2 will rapidly cluster, resulting in condensation of proteins with phase separation capacity. **(B)** Representative images of 293T cells transiently transfected with CRY2-mCh control, CRY2-mCh-CELF2, or CRY2-mCh-CELF2IDR constructs before and after blue light illumination. CRY2-mCh-CELF2 and CRY2-mCh-CELF2IDR formed optoDroplets, suggesting phase separation capacity of CELF2 IDR. Scale bar: 5 µm. **(C)** Quantification of optodroplets. Area and number of optodroplets in 3 representative cells were measured using FIJI. Statistical analysis: one-way ANOVA, *p < 0.05; ***p < 0.001; ****p < 0.0001. **(D)** Representative images of droplets formed by recombinant GFP-CELF2IDR in the droplet formation buffer containing 10% PEG-8000 at indicated protein concentrations. **(E)** Quantification of GFP-CELF2IDR droplets. Droplet size and number was measured using FIJI. **(F)** Schematic illustration of the LacO array system to test CELF2 self-interactions. eYFP-CELF2-LacI is recruited to LacO array through protein-DNA binding and mCherry-CELF2 can be recruited to the LacO array through CELF2-CELF2 interactions. **(G-H)** Quantification and representative images of mCherry-CELF2 recruitment to the LacO hub through CELF2-CELF2 self-association. mCherry-CELF2 signal is enriched at the LacO hub through interacting with eYFP-CELF2-LacI, while reduced or no mCherry enrichment was detected at the eYFP hub in the CELF2ΔIDR or mCherry control groups. Enrichment of mCherry above relative level of 1 suggests protein-protein interaction. Statistics: one-way ANOVA, **P < 0.01; ****P < 0.0001. Scale bar: 10 µm.

We further examined CELF2 condensation/self-interaction using a previously established imaging-based approach ^44–47^. Through specific LacI-LacO interaction, eYFP-LacI-tagged protein is recruited to the LacO array, generating a concentrated interaction hub. mCherry-tagged protein is brought to the hub through multivalent interactions and the intensity of mCherry at the eYFP hub can be measured to quantify the potency of these multivalent interactions (**Fig. 2F**). We co-expressed mCherry-CELF2 and eYFP-CELF2-LacI in U2OS cells containing a synthetic Lac operator (LacO) array integrated to the genome ^48^. We observed a strong mCherry signal at the hub (**Fig. 2G-H**), indicating a strong self-interaction ability of CELF2. When we performed the assay using IDR-truncated CELF2 (ΔIDR), we observed a much weaker mCherry signal at the array, even though the recruitment of eYFP-CELF2ΔIDR-LacI molecules to LacO array appeared normal (**Fig. 2G-H**). Therefore, these data suggest that CELF2 IDR is required for CELF2 condensation/self-interaction.

### CELF2 IDR is required for its function in regulating alternative splicing of tau and can be functionally substituted by FUS or TAF15 IDR

To understand the importance of CELF2 condensation on its function, we transfected GFP-tagged CELF2wt or CELF2ΔIDR in SH-SY5Y neuroblastoma cells and examined condensate formation and tau splicing. We observed that GFP-CELF2wt was localized to the nucleus and displayed a punctate distribution pattern (**Fig. 3A**), suggesting that CELF2 condensation might be associated with its function in the nucleus. In contrast, GFP-CELF2ΔIDR showed a homogeneous expression pattern in the nucleus, as reflected by the low fringe visibility measuring the GFP signals (**Fig. 3A, 3B**), indicating that the disordered hinged domain is required for CELF2 condensate formation. When we examined tau splicing, we found that SH-SY5Y cells transfected with the vector control expressed predominantly 3R tau. In cells transfected with GFP-CELF2wt, 4R tau expression was significantly enhanced (**Fig. 3C, 3D**), consistent with the data obtained from the Celf2 KO mouse brain showing that CELF2 functions to promote tau exon 10 inclusion. Unlike CELF2wt, CELF2ΔIDR failed to enhance 4R tau expression (**Fig. 3C, 3D**), indicating that CELF2 IDR is critical for its function in regulating alternative splicing.

**Fig. 3.**
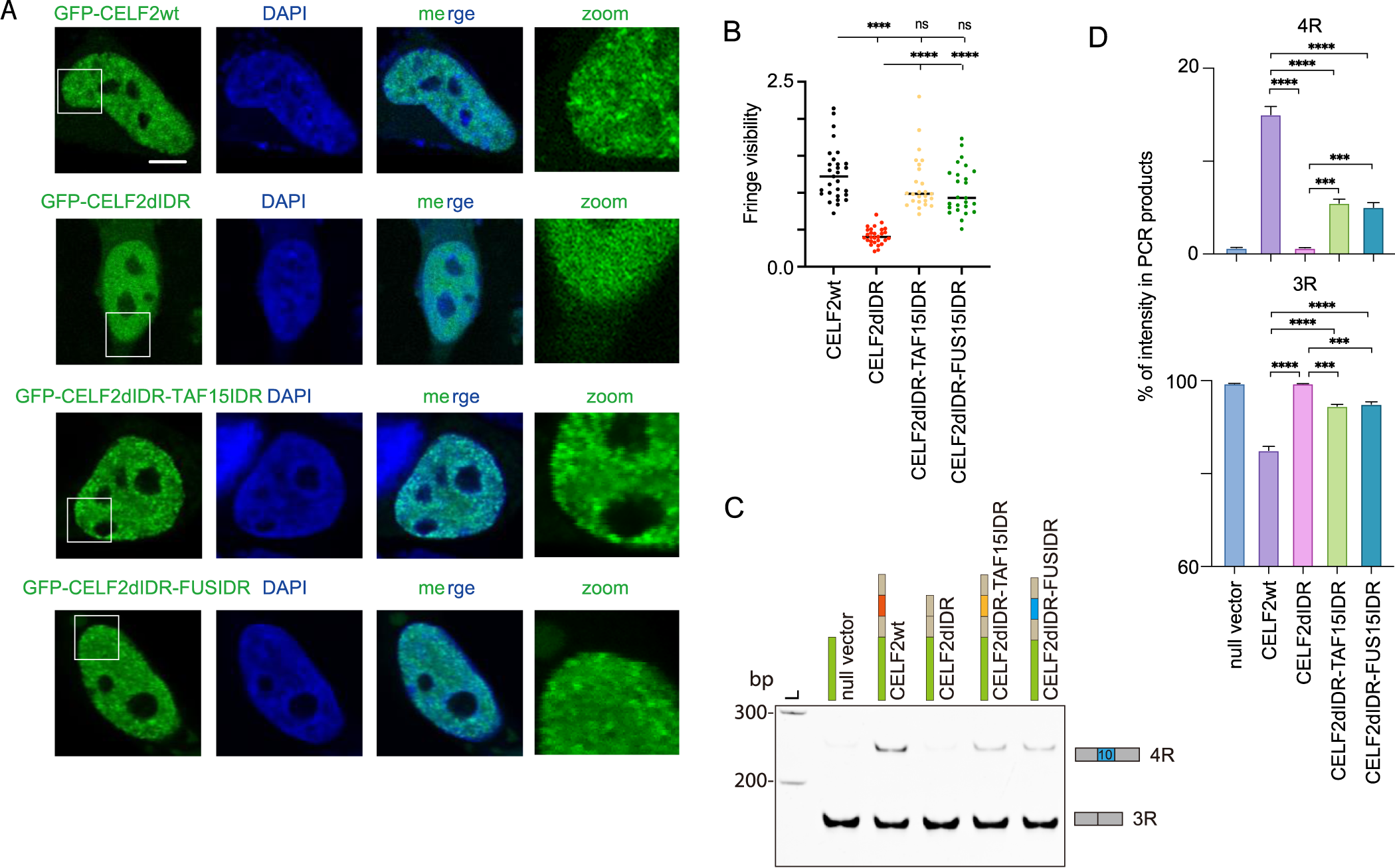
CELF2 IDR is required for its function in regulating alternative splicing of tau and can be functionally substituted by FUS or TAF15 IDR. **(A)** Representative confocal images of SH-SY5Y cells transiently transfected with the indicated CELF2 constructs. Deletion of CELF2IDR abolished CELF2 condensation, but replacing CELF2IDR with FUSIDR or TAF15IDR retained condensate formation capacity of CELF2. Scale bar: 5 µm. **(B)** Quantifications of fringe visibility of CELF2 condensates. Fringe visibility values of randomly selected AR foci from 6 cells (5 foci/cell) were plotted for each condition. Statistics: one-way ANOVA, ****P < 0.0001. **(C)** Representative gel image of RT-PCR results showing 4R and 3R tau level in SH-SY5Y cells transfected with the indicated constructs. Expression of CELF2wt but not CELF2ΔIDR promoted 4R tau expression, and CELF2IDR could be functionally replaced by the IDR of FUS or TAF15. **(D)** Quantification of RT-PCR results from (C). The intensity of 4R and 3R *MAPT* bands were measured using FIJI and the percentage of each band was calculated and plotted. Statistics: one-way ANOVA, ***p < 0.001; ****p < 0.0001.

It has been previous shown that in some cases one IDR can be functionally replaced by another in RNP granule assembly ^49,50^. We then asked whether CELF2 IDR could be substituted by other IDRs to regulate CELF2 condensate formation and function. Fused in sarcoma/translocated in liposarcoma (FUS/TLS or FUS) and TAF15 represent two of the most studied examples of proteins that undergo multivalent interactions and phase separation ^51^. The prion-like domain (PrLD) in FUS protein is an IDR enriched with tyrosine residues and can undergo phase separation driven by tyrosine-tyrosine interactions ^12^. TAF15 IDR has the same number of tyrosine residues but more charged residues than FUS IDR and exhibits a strong tendency to phase separate ^52^. We proceeded to replace CELF2’s IDR with IDRs from FUS and TAF15, inserted between RRM2 and RRM3 (**Fig. S3A, B**). Notably, the fusion proteins (CELF2ΔIDR with FUS or TAF15 IDR) display similar condensates in cells as CELF2wt (**Fig. 3A, 3B**). Furthermore, CELF2 with replaced IDR was sufficient to promote 4R tau expression (**Fig. 3C, 3D**). These results indicate that CELF2 IDR can be functionally substituted by selective IDRs. The correlation of condensate formation status and function in promoting 4R tau expression further support that the CELF2 condensation is critical for its function.

### CELF2 forms heterotypic condensates with NOVA2 and SFPQ through its IDR

Given that an alternative splicing (AS) event is regulated by multiple RNA-binding proteins (RBPs) and that IDRs facilitate both homotypic and heterotypic interactions, we sought to identify the RBPs that interact with CELF2, forming heterotypic condensates and collectively regulating AS. To capture CELF2-interacting proteins, we used the proximity-dependent biotin identification TurboID technique ^53^ with CELF2 as the bait protein (**Fig. 4A**). Fusing TurboID to CELF2 allowed labeling molecules interacting with CELF2. The labeled interactors were then pulled down with streptavidin peroxidase beads followed by proteomic profiling (**Fig. S4A**). Consistent with the function of CELF2 in regulating alternative splicing, the identified CELF2-interacting proteins were most significantly enriched for proteins involved in RNA splicing via spliceosome (**Fig. 4B**). We also performed TurboID for CELF2ΔIDR and found that IDR deletion diminished the interaction between CELF2 and a list of its interactors. KEGG pathway enrichment analysis on these interactors again identified RNA splicing-related proteins (**Fig. 4C**). Among the splicing-related proteins whose interaction with CELF2 was weakened by IDR deletion (**Fig. 4D**), we selected NOVA2 and SFPQ for further investigation due to their function in neuronal splicing and neurodegeneration ^54–56^.

**Fig. 4.**
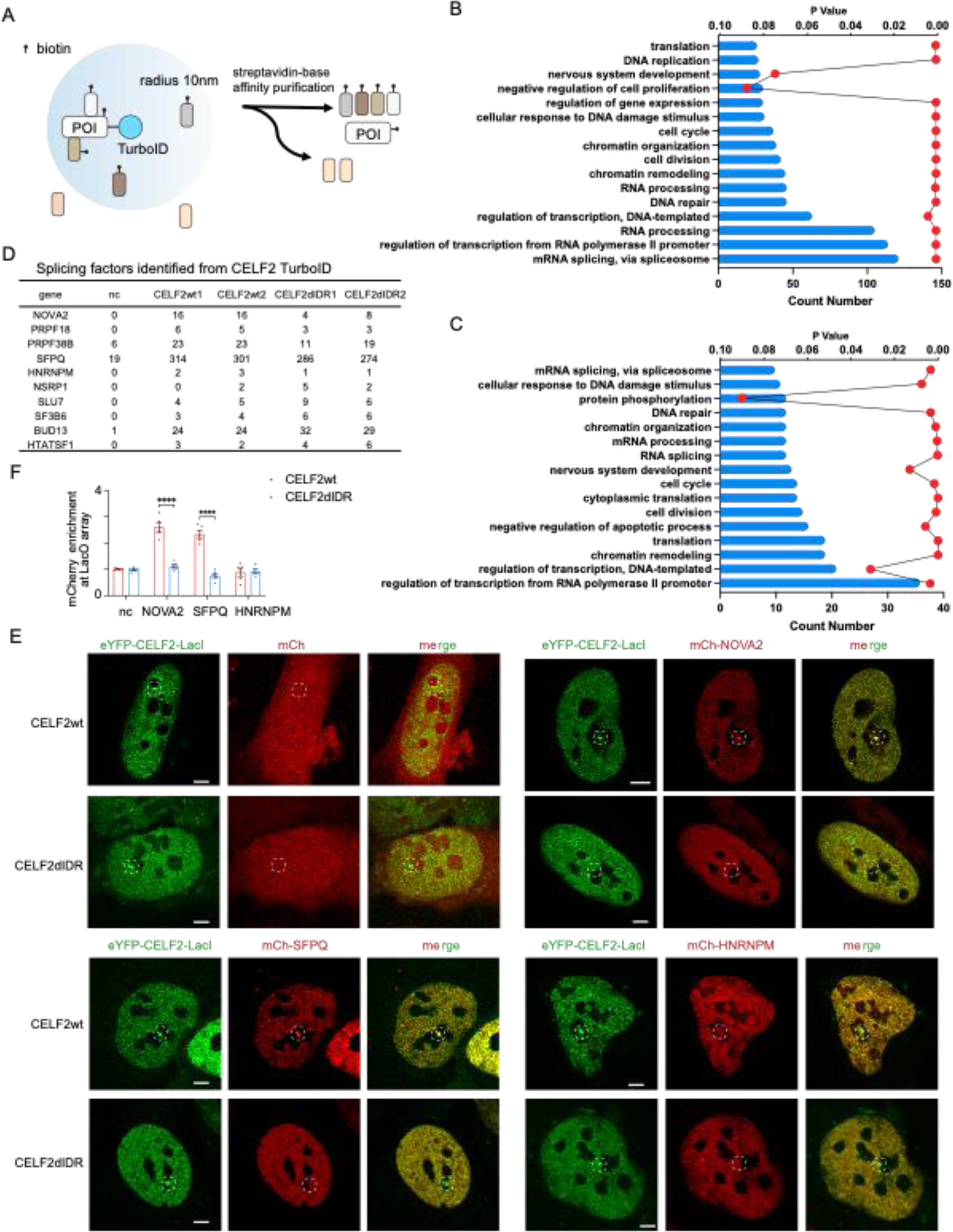
TurboID identified CELF2 cofactors with which CELF2 forms heterotypic condensates through its IDR. **(A)** Schematic illustration of proximity-dependent biotin identification technique, TurboID, to identify cofactors of CELF2wt and CELF2ΔIDR. **(B)** KEGG pathway enrichment analysis on turboID-identified CELF2 cofactors. Top 16 pathways ranked by Count are plotted. mRNA splicing pathway is among the most significantly enriched pathways. **(C)** KEGG pathway enrichment analysis on turboID-identified CELF2 cofactors that show differential interaction with CELF2wt vs CELF2ΔIDR. Top 16 pathways ranked by Count are plotted. **(D)** Peptide numbers of the splicing factors identified from Turbo ID using empty vector, CELF2wt or CELF2ΔIDR as baits. Peptide numbers of two independent experiments are shown. NOVA2 and SFPQ showed reduced peptide numbers in CELF2ΔIDR than in CELF2wt TurboID, and were then selected for further experiments to test their interactions with CELF2. **(E)** Representative images of mCh-NOVA2, mCh-SFPQ and mCh-HNRNPM recruitment to the LacO hub through CELF2-cofactor interactions in U2OS cells. mCh-NOVA2 and mCh-SFPQ, but not mCh-HNRNPM, were enriched at the LacO hub, and the enrichment was dependent on CELF2 IDR. Scale bar: 5 µm **(F)** Quantification of the multivalent interactions between CELF2 and its cofactors. Enrichment of mCherry above relative level of 1 suggests protein-protein interaction. Statistics: one-way ANOVA, ****P < 0.0001.

We next interrogated whether CELF2 co-localized with SFPQ and NOVA2 in cells. We selected 293T cells for our study due to their low expression levels of CELF2, ensuring that the introduced expression patterns would not be confounded by endogenous CELF2 expression. When expressed individually, mCh-NOVA2 and mCh-SFPQ were localized in the nucleus, with a weakly punctate pattern (**Fig. S4B**). When co-expressed with GFP-CELF2, we observed a high level of co-localization between GFP-CELF2 and mCh-NOVA2 (or mCh-SFPQ) (**Fig. S4C, S4D**). Interestingly, in the presence of GFP-CELF2, both mCh-NOVA2 and mCh-SFPQ displayed a strongly punctate pattern as measured by the higher fringe visibility of mCherry signal (**Fig. S4C-F**). This suggests that CELF2 might facilitate to recruit NOVA2 and SFPQ to those nuclear condensates. We further found that the CELF2-dependent NOVA2 and SFPQ condensates required the IDR of CELF2, as mCh-NOVA2 and mCh-SFPQ showed significantly weaker condensation when co-expressed with CELF2ΔIDR (**Fig. S4C-F**).

To further investigate the heterotypic interactions between CELF2 and cofactors, we employed the LacO array imaging assay. We co-expressed eYFP-CELF2-LacI and mCherry-tagged cofactors in U2OS cells containing a synthetic Lac operator (LacO) array. Compared to mCherry control which did not show enrichment at the eYFP+ hub, mCh-NOVA2 and mCh-SFPQ was highly enriched (**Fig. 4E, 4F**), suggesting a strong multivalent interaction between CELF2 and NOVA2 or SFPQ. As a negative control, we also examined HNRNPM, which was identified as a weak CELF2 interactor from TurboID (**Fig. 4D**), and observed no enrichment of mCh-HNRNPM on the eYFP+ hub, indicating the specificity of CELF2-NOVA2 and CELF2-SFPQ interactions. Notably, these heterotypic interactions were dependent on CELF2 IDR, as IDR deletion abolished the enrichment of NOVA2 and SFPQ (**Fig. 4E, 4F**). Together, these data suggest that CELF2 forms heterotypic condensates with NOVA2 and SFPQ through its IDR.

### CELF2 co-regulates tau splicing with NOVA2 and SFPQ

Having shown that CELF2 interacts with NOVA2 and SFPQ in nuclear condensates, we asked if they functioned together to regulate AS. Similar to CELF2, overexpression of SFPQ in SH-SY5Y cells led to enhanced 4R tau expression, although its effect was not as strong as CELF2 (**Fig. 5A, 5B**). Simultaneously expressing both CELF2 and SFPQ resulted in a higher 4R tau level than expressing either CELF2 or SFPQ alone (**Fig. 5A, 5B**), while co-expression of SFPQ and CELF2ΔIDR had a similar effect as SFPQ overexpression. Therefore, both CELF2 and SFPQ can promote tau exon 10 inclusion and they can coordinate to achieve a higher level of 4R tau expression.

**Fig. 5.**
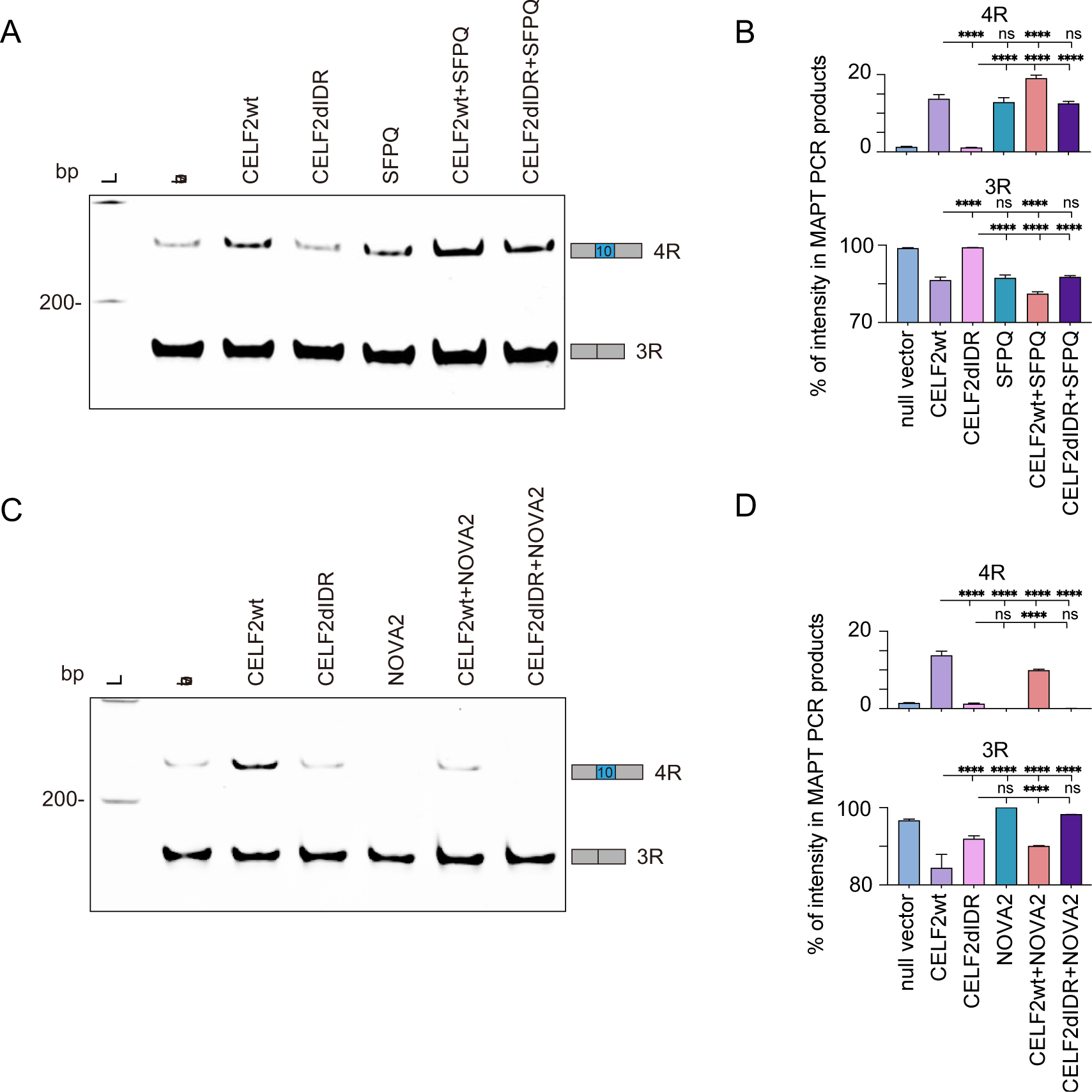
CELF2 co-regulates tau splicing with NOVA2 and SFPQ. **(A)** Representative gel image showing 4R and 3R tau expression in SH-SY5Y cells transfected with. with indicated plasmid(s). Cells were transfected for 48 hours before RNA extraction and RT-PCR to examine alternative splicing of the endogenous *MAPT* gene. **(B)** Quantification of RT-PCR shown in (A). Both CELF2 and SFPQ promote 4R tau expression. Co-expression CELF2 and SFPQ had a greater effect on exon 10 inclusion than either CELF2 or SFPQ expression alone. FIJI was used to measure the intensity of each *MAPT* transcript product and the percentages of 4R and 3R tau were plotted. Statistics: one-way ANOVA, ****, p < 0.0001. **(C)** Representative gel image showing the effect of CELF2 and NOVA2 on *MAPT* alternative splicing. SH-SY5Y cells were transfected with indicated plasmid(s) for 48 hours, followed by RNA extraction and RT-PCR. **(D)** Quantification of RT-PCR shown in (C). CELF2 promotes exon 10 inclusion, while NOVA2 promotes exon 10 skipping. Co-expression of NOVA2 with CELF2 dampened the effect of CELF2 on promoting 4R tau expression. FIJI was used to measure the intensity of each *MAPT* transcript product and the percentage of 4R and 3R tau was plotted. Statistics: one-way ANOVA, ****p < 0.0001.

Unlike CELF2 or SFPQ, NOVA2 overexpression inhibited tau exon 10 inclusion, resulting in a complete depletion of 4R tau (**Fig. 5C, 5D**). In addition, NOVA2 expression was sufficient to diminish CELF2-induced 4R tau expression, suggesting that NOVA2 might compete with CELF2 to promote exon 10 skipping. To further understand the function of NOVA2 in regulating tau splicing, we generated stable SH-SY5Y cell lines expressing Dox-inducibe shRNAs targeting NOVA2 (**Fig. S5A**). We noticed that 4R tau expression was barely detectable in the stable cell line. This was likely due to maturation of SH-SY5Y cells during the process of establishing stable cell lines (**Fig. S5B, S5C**). Upon Dox treatment to induce NOVA2 knockdown, 4R tau expression was significantly elevated (**Fig. S5B, S5C**), again suggesting that NOVA2 function to inhibit exon 10 inclusion. We next established stable cell lines with Dox-inducible CELF2 expression and shRNA targeting NOVA2 to test if CELF2-induced exon 10 inclusion could be elevated after knocking down NOVA2. Indeed, we found that knocking down NOVA2 in cells with CELF2 overexpression was able to further enhance 4R tau expression (**Fig. S5D, S5E**). Together, these data suggest that CELF2 interacts with other RBPs in biomolecule condensates to co-regulate AS, and that the components within the condensates determine the splicing outcomes.

### The charged residue D388 is critical for CELF2 condensate properties and CELF2 function

IDRs that mediate phase separation (PS) are often enriched with charged amino acids, polar amino acids, and/or aromatic amino acids, which have all been proposed to contribute to IDR PS ^11–17^. In the *in vitro* droplet formation assay, we observed that the droplet size and number negatively correlated with the concentration of NaCl in the droplet formation buffer (**Fig. 6A, 6B)**. Based on their overall charge composition, IDRs can exhibit either salt-out or salt-in behavior. This phenomenon is influenced by the screening of electrostatic interactions and Hofmeister effects ^57–59^. The sensitivity of IDR droplets to salt concentration suggest that charged residues might contribute to CELF2 phase separation. Within CELF2 IDR, there is only one negatively charged residue (D388) and zero aromatic residue (**Fig. S6A**). Intriguingly, this negatively charged amino acid is conserved among all 6 CELF proteins (**Fig. S6B**), suggesting its functional importance.

**Fig. 6.**
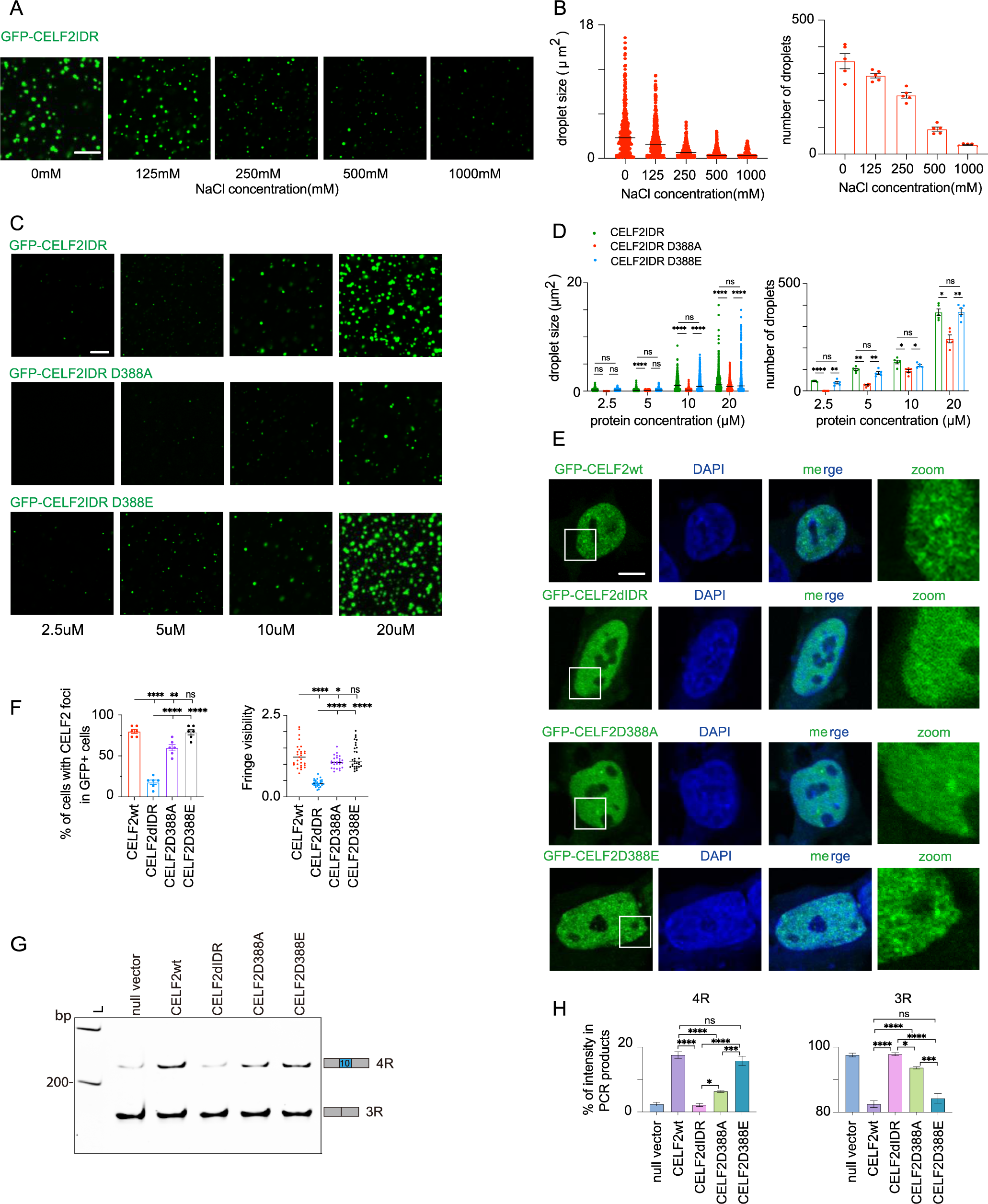
The charged residue D388 is critical for CELF2 condensate properties and CELF2 function. **(A)** Representative images of *in vitro* droplets formed by recombinant CELF2 IDR in droplet formation buffers containing10% PEG-8000 and with various salt concentrations. Scale bar: 5 µm. **(B)** Quantification of CELF2IDR droplet size and number at indicated conditions using FIJI. Droplet size and number negatively correlates with NaCl concentration. **(C)** Representative images of *in vitro* droplets formed by wildtype and mutant CELF2 IDRs at indicated protein concentrations. Scale bar: 5 µm. **(D)** Quantification of droplet size and number of wildtype and mutant CELF2 IDRs at indicated protein concentrations. D388A mutation led to smaller and fewer droplets, while D388E mutation did not affect droplet size or number. **(E)** Representative confocal images of GFP-CELFwt, GFP-CELF2ΔIDR, GFP-CELFD388A and GFP-CELF2D388E in SH-SY5Y cells. Scale bar: 5 µm. **(F)** Quantification of CELF2 condensates in SH-SY5Y cells. D388A mutation resulted in an reduction in the percentage of cells with CELF2 condensates, as well as a lower fringe visibility of GFP+ condensates. Statistics: one-way ANOVA, *p < 0.05, **p < 0.01, ****p < 0.0001. **(G)** Representative RT-PCR gel image showing 4R and 3R tau expression in SH-SY5Y cells transfected with wildtype and mutant CELF2 genes. **(H)** Quantification of RT-PCR shown in (G). D388A mutation significantly reduced the effect of CELF2 in promoting 4R tau expression, while D388E did not show much impact. FIJI was used to measure the intensity of each *MAPT* transcript product and the percentages of 4R and 3R tau were plotted. Statistics: one-way ANOVA, *p < 0.05, ***p < 0.001, ****p < 0.0001.

To investigate whether the negative charge of D388 plays a role in regulating CELF2 condensation and function, we generated two CELF2 variants: D388A to replace the negatively charged D residue with a non-charged residue A, and D388E to replace the aspartic acid with another negatively charged amino acid glutamic acid. We first purified the IDR fragments (**Fig. S6C**) and performed *in vitro* droplet assay. We found that both GFP-CELF2IDR(D388A) and GFP-CELF2IDR(D388E) formed droplets. However, the D388A droplets were significantly smaller and fewer than WT when examined at the same protein concentrations (**Fig. 6C, 6D**). And the minimal concentration required for droplets formation was higher for to IDR(D388A), suggesting that the D to A mutation inhibits CELF2 IDR condensation propensity. Interestingly, GFP-CELF2IDR(D388E) behaved similarly as GFP-CELF2IDR(WT), forming comparable sizes and numbers of droplets at all concentrations tested (**Fig. 6C, 6D**).

We next determined whether D388 was required for CELF2 condensation in cells and for CELF2 function in regulating alternative splicing. In line with the results of *in vitro* droplet formation assay, GFP-CELF2(D388A) formed less distinct condensates in SH-SY5Y cells, as measured by the reduced fringe visibility (**Fig. 6E, 6F**). D388A also significantly reduced the number of cells with CELF2 condensates, although the effect was not as strong as ΔIDR. In contrast, D388E did not show significant changes in condensate fringe visibility or number of cells with condensates when compared to WT (**Fig. 6E, 6F**). Like CELF2(WT), which induced tau exon 10 inclusion upon overexpression in SH-SY5Y cells, both CELF2(D388A) and CELF2(D388E) were able to promote the expression of 4R tau. However, the expression level of 4R tau in cells expressing CELF2(D388A) was lower than that observed in cells with CELF2(WT) expression, whereas the level of 4R tau in CELF2(D388E) expressing cells was comparable to that in CELF2(WT) expressing cells (**Fig. 6G, 6H**). This was not due to a difference in the expression level of CELF2 variants (**Fig. S6D**). Therefore, replacing the negatively charged D388 with a non-charged residue abolished CELF2 condensation and function, while replacing D388 with another negatively charged amino acid did not affect CELF2 condensation or function.

As CELF2 requires its IDR to interact with NOVA2 an dSFPQ (**Fig. 4),** we asked whether D388 influenced CELF2’s heterotypic interactions using the LacO array imaging assay. We co-expressed eYFP-CELF2 (WT or D388A)-LacI and mCh-NOVA2 (or mCh-SFPQ) in U2OS 2-6-3 cells. The enrichment of mCh-NOVA2 or mCh-SFPQ at the eYFP+ hubs was significantly eliminated by D388A mutation (**Fig. 7A, 7B),** suggesting that the charged residue D388 is critical for CELF2 heterotypic interactions. Taken together, these data suggest that the negatively charged residue within CELF2 IDR can modulate CELF2 condensation and thus impact its function.

**Fig. 7.**
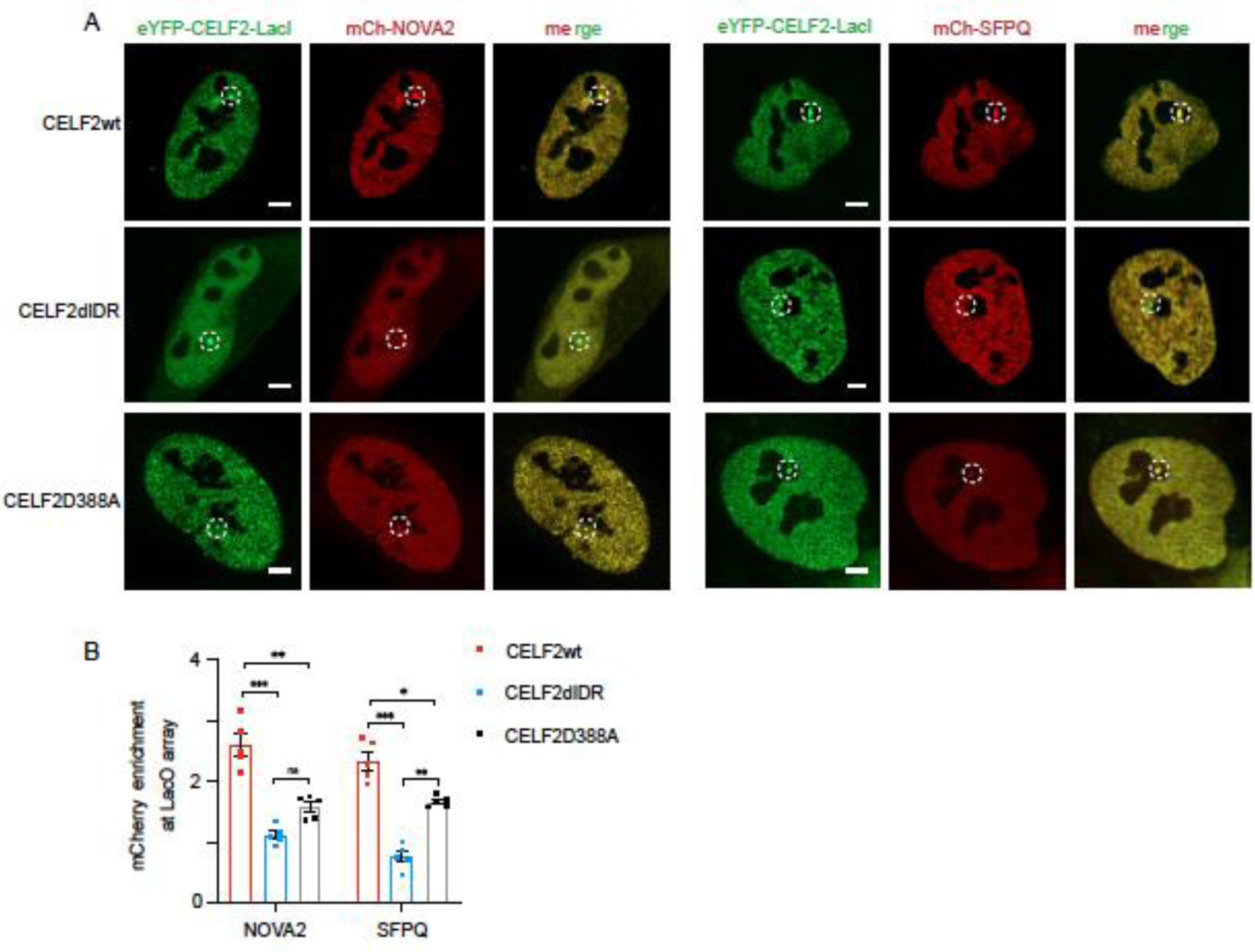
D388 within CELF2 IDR is critical for CELF2 interaction with NOVA2 and SFPQ. **(A)** Representative images of mCh-NOVA2 and mCh-SFPQ enrichment at the LacO hub. U2OS cells carrying the LacO array were co-transfected with EYFP-CELF2-LacI (*wt*, *ΔIDR*, or *D388A*) and a mcherry vector (mCh-NOVA2, mCh-SFPQ, or mCh-null vector). Scale bar: 5 µm **(B)** Quantification of the multivalent interactions between CELF2 and NOVA2 (or SFPQ). Enrichment of mCherry above relative level of 1 suggests protein-protein interaction. D388A mutation partially abolished CELF2-cofactor interactions. Statistics: one-way ANOVA, ****P < 0.0001.

## DISCUSSION

The orchestration of every alternative splicing event depends on the concerted effort of multiple RBPs. But how different proteins are recruited to the splicing sites are not clear. Through multiple approaches, we demonstrated that CELF2 forms phase-separated concentrates with other splicing regulators through its disordered hinge domain to regulate alternative splicing. Among the CELF2 targets identified by our CLIP-seq, we selected tau to study the functional significance of CELF2 condensation and its regulation. Our *in vivo* and *in vitro* studies revealed that CELF2 acts to promote tau exon 10 inclusion. The intrinsically disordered hinge domain of CELF2 mediates CELF2 condensation and activity, and can be functionally replaced by IDRs from FUS and TAF15. Our TurboID identified CELF2-interacting proteins including NOVA2 and SFPQ, which co-condensated with CELF2 to regulate alternative splicing, with NOVA2 inhibiting but SFPQ promoting tau exon 10 inclusion. The finding that the negatively charged aspartic acid residue within the IDR (D388) is essential for both homotypic and heterotypic interactions, as well as for CELF2’s function, aligns with the conservation of this critical residue among CELF paralogs. These data highlight the importance of the disordered hinge domain of RBPs in mediating the formation of biomolecular condensates and suggest that the makeup of proteins within the splicing condensates dictates the outcomes of alternative splicing events.

### CELF2 is involved in AD by regulating alternative splicing

Alternative splicing affects more than 95% of human genes, significantly contributing to the diversity and complexity of proteins ^60,61^. Recent investigations into alternative splicing in aging brains have uncovered specific aberrant splicing events linked to AD ^62–64^. Moreover, genome-wide association studies (GWAS) have identified associations between CELF genes and various inherited neurological conditions ^65,66^, and specific single nucleotide polymorphisms (SNPs) in the CELF2 gene have been linked to AD ^28,29^. While these studies provide important insights into potential genetic links, the exact role of CELF2 in the pathogenesis of Alzheimer’s disease remains unclear. Our CLIP-seq revealed that CELF2 targets are enriched with genes involved in neurodegenerative diseases. Loss of CELF2 consistently leads to a reduction in the 4R:3R ratio, even though this ratio in the mouse brain gradually changes during development. However, it remains to be investigated whether the dynamic changes in the 4R:3R ratio during development and aging are related to variations in CELF2 expression. Manipulation of this ratio has been demonstrated to induce AD-like phenotypes ^67,68^. Therefore, CELF2 may play a crucial role in the development of AD by controlling the levels of tau isoforms. Certainly, the function of CELF2 in AD is not limited to its regulation on Tau splicing, as many other AD-related genes are targets of CELF2 and CELF2 regulates many aspects of RNA metabolism and function.

### The divergent domain regulates CELF2 function and can be substituted by other IDRs

All six CELF members share a similar structure, which includes three RRMs, two located at the N-terminal and one at the C-terminal, with a divergent domain situated between RRM2 and RRM3. The divergent domain exhibits limited homology among CELF protein family members and does not show significant homology to other proteins. RRMs of CELF proteins are known to play important roles in mediating protein-RNA interactions ^19^. While the divergent domain itself may not function as an RNA-binding domain, evidence suggests it likely plays a crucial role in facilitating RNA interaction. In previous yeast three-hybrid experiments, deletions within the divergent domain had the most significant impact on CELF1 binding, surpassing alterations in the RRMs ^69,70^. Intriguingly, studies utilizing chimeric proteins revealed that the sequence of the divergent domain associated with the first two RRMs of CELF4 strongly influences binding ^71^. However, it remains unclear whether the divergent domain affects RNA binding through mediating protein-protein interactions or by conveying essential conformational states. In this work we show that the divergent domain is sufficient to form phase-separated droplets *in vitro* and is required for CELF2 condensation in cells. Consistent with previous studies, deletion of the divergent domain leads to a significant functional impairment (**Fig. 3**). Interestingly, CELF2 IDR (divergent domain) can be functionally substituted by FUS and TAF15 IDRs (Figure 3). FUS/TAF15 IDRs can self-associate to form phase-separated condensates under physiological conditions ^72^. The functional substitution indicate that the specific sequence of the divergent domain may not be critical for CELF2’s function. Rather, the intrinsically disordered nature of this domain plays a pivotal role in the formation of membraneless condensates. This mechanism has been shown to be essential for recruiting various components and enhancing local concentrations to facilitate functional activities ^40^. It’s worth noting that although chimeric proteins with FUS/TAF15 IDR and CELF2ΔIDR formed condensates in cells and promoted 4R tau splicing, they were not as effective as CELF2 in enhancing 4R tau splicing. This partial restoration could be attributed to the unique biophysical characteristics of condensates formed by CELF2 IDR compared to those formed by other IDRs. This observation aligns with our previous research, which demonstrated the influence of condensate biophysical properties on protein function ^47^.

### CELF2 and other splicing factors form dynamic splicing condensates to co-regulate alternative splicing

Within the genome, the same genomic sequences often serve as binding sites for multiple RBPs. RBPs frequently collaborate in complexes during splicing, influencing each other’s binding specificity and functional outcomes. This interplay among RBPs adds layers of complexity to the regulatory landscape of RNA processing and gene expression. Through turboID, we identified CELF2-interacting proteins, particularly proteins interacting with CELF2 through CELF2 IDR. Among them, we focused on NOVA2 and SFPQ due to their known function in regulating alternative splicing and association with neurodegenerative diseases. Cellular localization of these proteins suggests that CELF2 might recruit NOVA2 and SFPQ to nuclear condensates through IDR-mediated heterotypic interactions between CELF2 and cofactors (**Fig. 4**). Functionally, both CELF2 and SFPQ can promote tau exon 10 inclusion and they can coordinate to achieve a higher level of 4R tau expression. In contrast, NOVA2 acts to inhibit tau exon 10 inclusion and might compete with CELF2 (**Fig. 5**). These findings support that CELF2 interacts with other splicing regulators within biomolecule condensates, jointly influencing AS. Moreover, they suggest that the composition of these condensates determines the splicing outcomes. The balance between 4R and 3R tau isoforms in the human brain changes throughout development and aging, with 3R tau predominantly prevailing during brain development, and a gradual increase in 4R tau levels occurring, eventually reaching parity with 3R tau in adulthood ^73^. Thus, we propose that the splicing condensates undergo dynamic alterations to achieve temporally and spatially specific alternative splicing.

### The conserved negatively charged residue D388 modulates CELF2 condensation

A wide range of molecular forces contribute to the formation of biomolecular condensates ^11–17^. IDRs are commonly enriched with aromatic residues (phenylalanine, tryptophan and tyrosine) which mediate pi-pi interactions. Other interactions that have been suggested as important drivers of biomolecular phase separation are charge-charge interactions and Pi-cation interactions. In addition, hydrophobic amino acids, such as leucine, isoleucine, and valine, are often enriched in protein regions that drive phase separation. While hydrophobic interactions may be less predominant in the context of phase separation compared to folded proteins, they likely remain important due to the significant presence of hydrophobic amino acids ^74^. There is only 1 charged residue (D388) and no aromatic residue within the divergent domain. The negatively charged residue is conserved among the 6 CELF family members. We found that replacing D388 with an un-charged amino acid (Alanine) reduced CELF2 IDR droplet formation and tau exon 10 inclusion. D388A also diminished the interaction between CELF2 and NOVA2 or SFPQ (**Fig. 7**). Although the charged D388 plays an important role in modulating CELF2 condensation and function, it might not be the driving force for CELF2 IDR condensation. This is because charge-driven phase separation often occur in IDRs containing both positively and negatively charged residues or between two oppositely charged biomolecules ^74^. Therefore, the hydrophobic amino acids in IDR might be the major driving force for CELF2 condensation given their high abundance. Our observations that mutation of a conserved residue within CELF2 IDR was sufficient to alter CELF2 condensation and function in alternative splicing suggest that targeting CELF2 might represent a potential strategy to intervene neurodegenerative diseases.

### Limitations of the study

Splicing reactions occur within the spliceosome, a dynamic RNA-protein complex. This complex exhibits remarkable plasticity in substrate recognition, allowing it to incorporate various segments of a pre-mRNA into a mature mRNA, under the influence of numerous regulatory proteins ^2^. RNAs can orchestrate the assembly of multiple RBPs to condensates and dictate condensate composition ^75^. Furthermore, RNA secondary structure can influence the physical characteristics of a condensate and, independently of proteins, facilitate the formation of gel-like phases. Moreover, RNAs can modify protein conformation to trigger phase separation in response to environmental cues ^76–78^. Our study reveals that CELF2 and other RBPs co-condensate to regulate mRNA alternative splicing. However, it remains to be investigated how mRNA impact the physical properties of CELF2 condensates.

## Acknowledgements

We would like to thank David Libich for suggestions, and all members in Chen and Liu labs for technical assistance and helpful discussions. This work was supported by funds from Voelcker Fund Young Investigator Award to L.C., NIA R01AG070214 to L.C., NIA R01AG071591 to L.C., CPRIT RR160017 to Z.L., V Foundation V2016-017 to Z.L., V Foundation DVP2019-018 to Z.L., Voelcker Fund Young Investigator Award to Z.L., UT Rising STARs Award to Z.L., Susan G. Komen CCR Award CCR17483391 to Z.L., NCI U54 CA217297/PRJ001 to Z.L., the Mary Kay Foundation Cancer Research Grant to Z.L., and NIGMS R01GM137009 to Z. L..

## Author Contributions

L.C., and Z.L. conceived the work and designed the study. X.L., I.S., R.A., D.T., and S.K. performed experiments and data analyses. Z.Z. performed the computational analyses for CLIP-seq and Turbo ID data. L.C. supervised the research and oversaw the project. L.C. Z.L. and X.L. wrote the manuscript with input from all authors.

## Resource Availability

### Lead Contact

Further information and requests for resources and reagents should be directed to and will be fulfilled by the lead contact, Lizhen Chen

### Materials availability

All unique materials generated in this study, including strains and plasmids will be shared by the lead contact upon request.

### Data and code availability

Key quantitative data will be supplied in a source data file. Key images will be deposited in the Figshare repository. Additional data reported in this paper are available from the lead contact upon request.

