## supplemental figures for "CELF2 promotes tau exon 10 inclusion via hinge domain-mediated nuclear condensation"

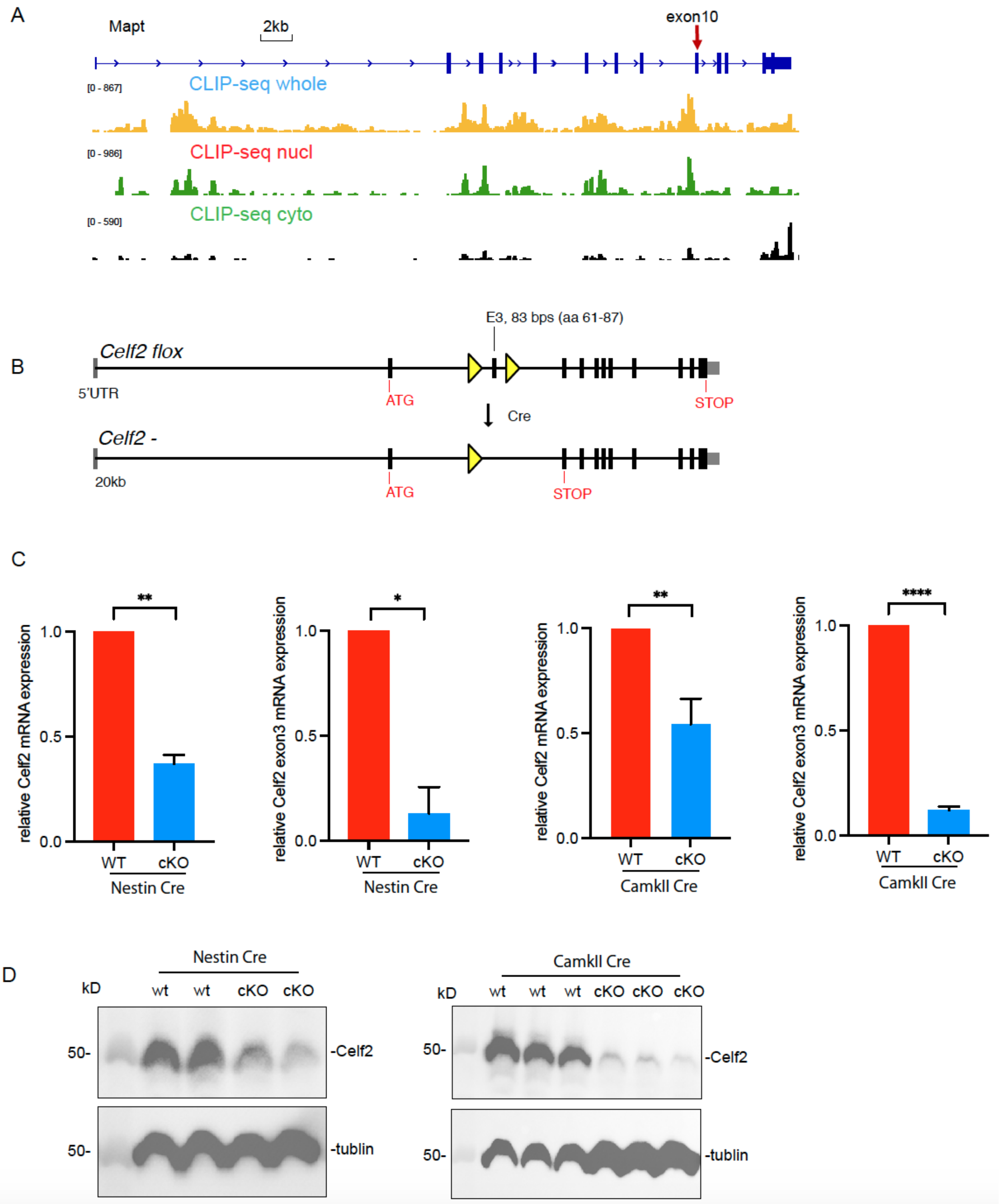


**Fig. S1. The binding sites of CELF2 on *Mapt* mRNA and validation of Celf2 cKO mouse (related to Fig. 1).**

**(A)** Genome browser view of CLIP-seq peaks on the *Mapt* gene locus. CLIP-seq was performed using mouse N2A cells and the CLIP-seq reads were aligned to mouse genome (mm9). CLIP-seq results from whole cell lysate, cytoplasmic fraction, and nuclear fraction show that CELF2 binds to 3'-UTR of Mapt mRNA in the cytoplasm, and binds to the intron 5'-to the alternatively spliced exon 10 in the nucleus.

(B) Schematic illustration showing the *Celf2* cKO allele (flox allele). Exon 3 is flanked by two *flox* elements, allowing for exon 3 deletion upon *Cre* expression.

(C) RT-qPCR showing that *Celf2* cKO brains have reduced Celf2 mRNA expression. qRT-PCR was done using hippocampal and cortical tissues of *Nest-CreER;Celf2*$^{flox/flox}$ (4 weeks old), *CamKII-CreER;Celf2*$^{flox/flox}$ (2 months old), and littermate control mice. Two sets of PCR primers were used, one set to amplify 3' end of Celf2 mRNA and the other set to amplify exon3 specifically. Celf2 transcript was detected in cKO brains as Celf2 is only deleted in some cells. to perform RT-qPCR and measure total *Celf2* mRNA and *Celf2* mRNA containing exon 3.

(D) Western blot showing that *Celf2* cKO brains have reduced Celf2 protein expression. Western blot was done using hippocampal and cortical tissues of *Nest-CreER;Celf2*$^{flox/flox}$ (4 weeks old), *CamKII-CreER;Celf2*$^{flox/flox}$ (2 months old), and littermate control mice.

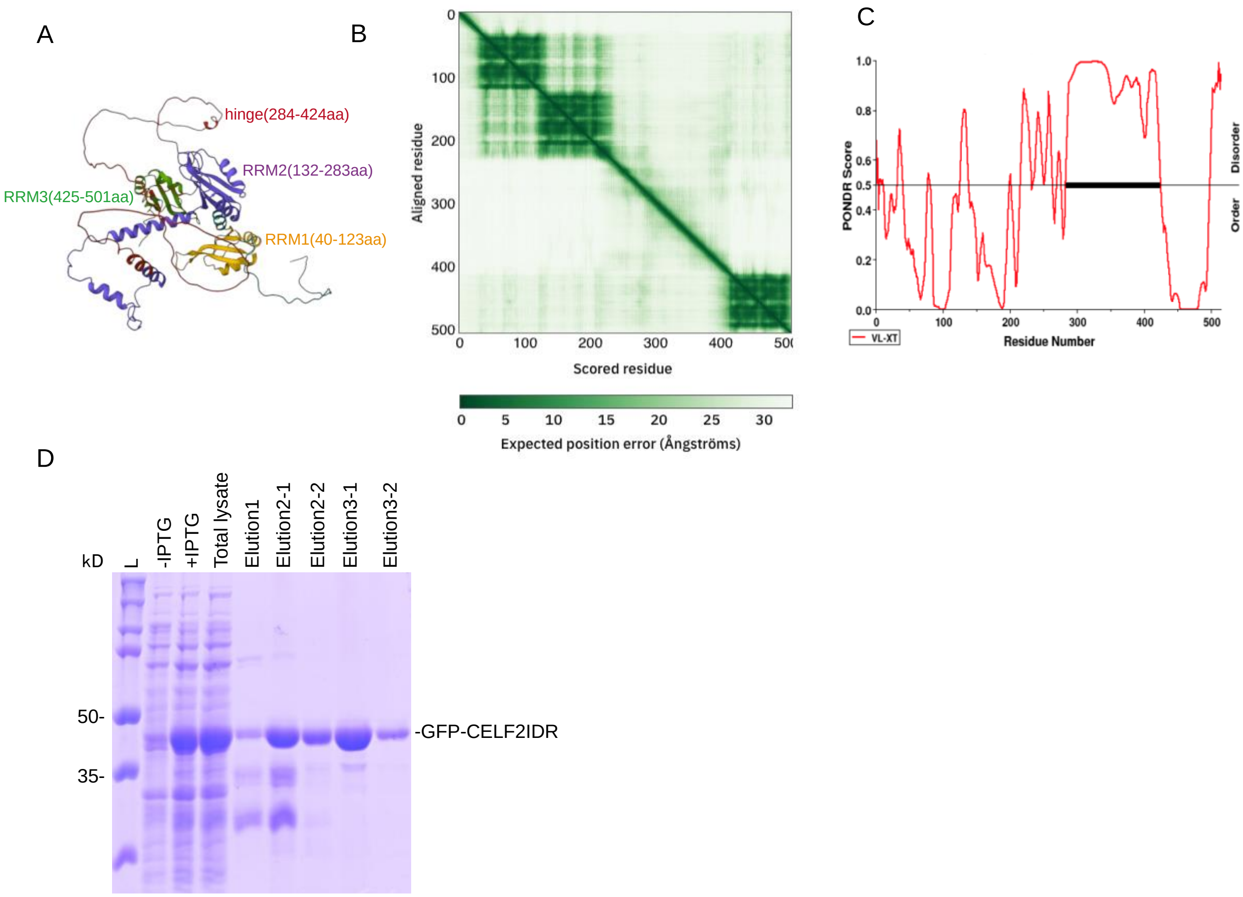


**Fig. S2. CELF2 hinge domain is a low complexity domain (related to Fig. 2).**

**(A)** Structure prediction of full-length CELF2 protein by AlphaFold (https://alphafold.ebi.ac.uk/). The hinge domain (284- 424) does not exhibit a defined structure.

**(B)** Predicted aligned error (PAE) plot for CELF2 protein indicating inter-domain accuracy. The shade of green shows expected distance error, with dark green indicating low error and light green indicating high error.

**(C)** Plot of intrinsic protein disorder probability for human CELF2 protein using PrDos (http://prdos.hgc.jp/cgi-bin/top.cgi) showing that the hinge domain has high disorder score.

**(D)** Coomassie staining of GFP-CELF2IDR (hinge domain) from indicated protein samples. Prior to protein purification, we compared cell lysates from cells without and with IPTG induction. We observed an induced band between 35 KDa and 50 KDa, which matched the size of His-GFP-CELF2NTD (43 KDa). Total lysate from protein purification experiment, flow through and elutes from Ni+ columns were also examined.

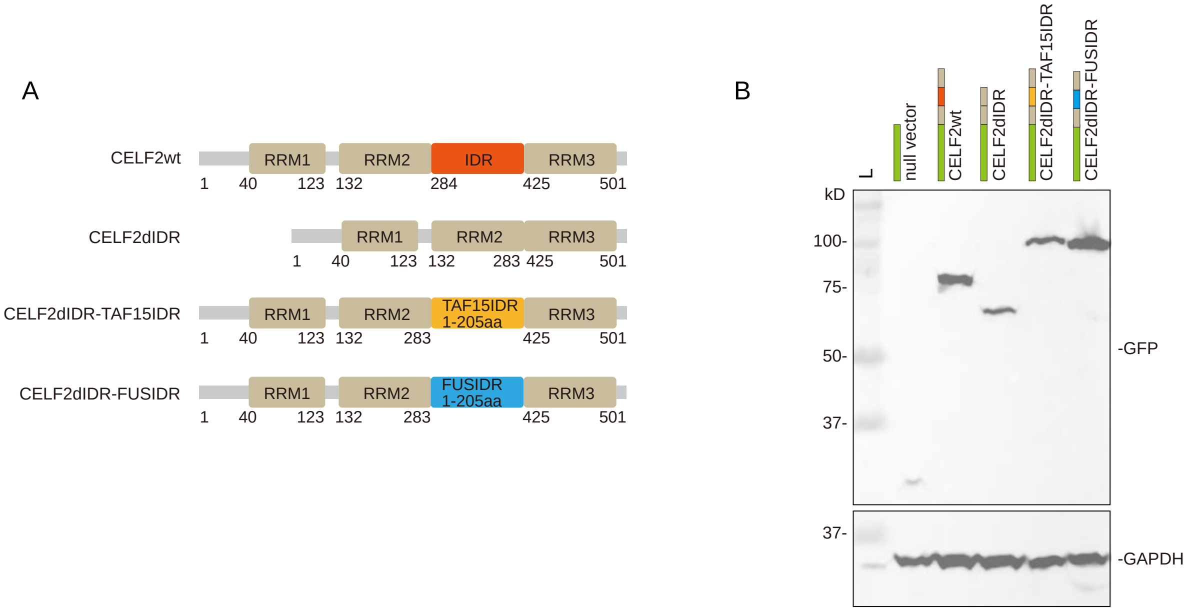


**Fig. S3. Replacing CELF2 IDR with IDRs of TAF15 and FUS (related to Fig. 3).**

**(A)** Schematic diagram of the GFP-CELF2 mutant fusion protein constructs.
**(B)** Indicated deletion and fusion proteins were expressed in cells to confirm that their expression levels are comparable. Cells were harvested 48 hours post-transfection for total protein extraction, followed by Western blotting using anti-GFP antibodies to detect their expression.

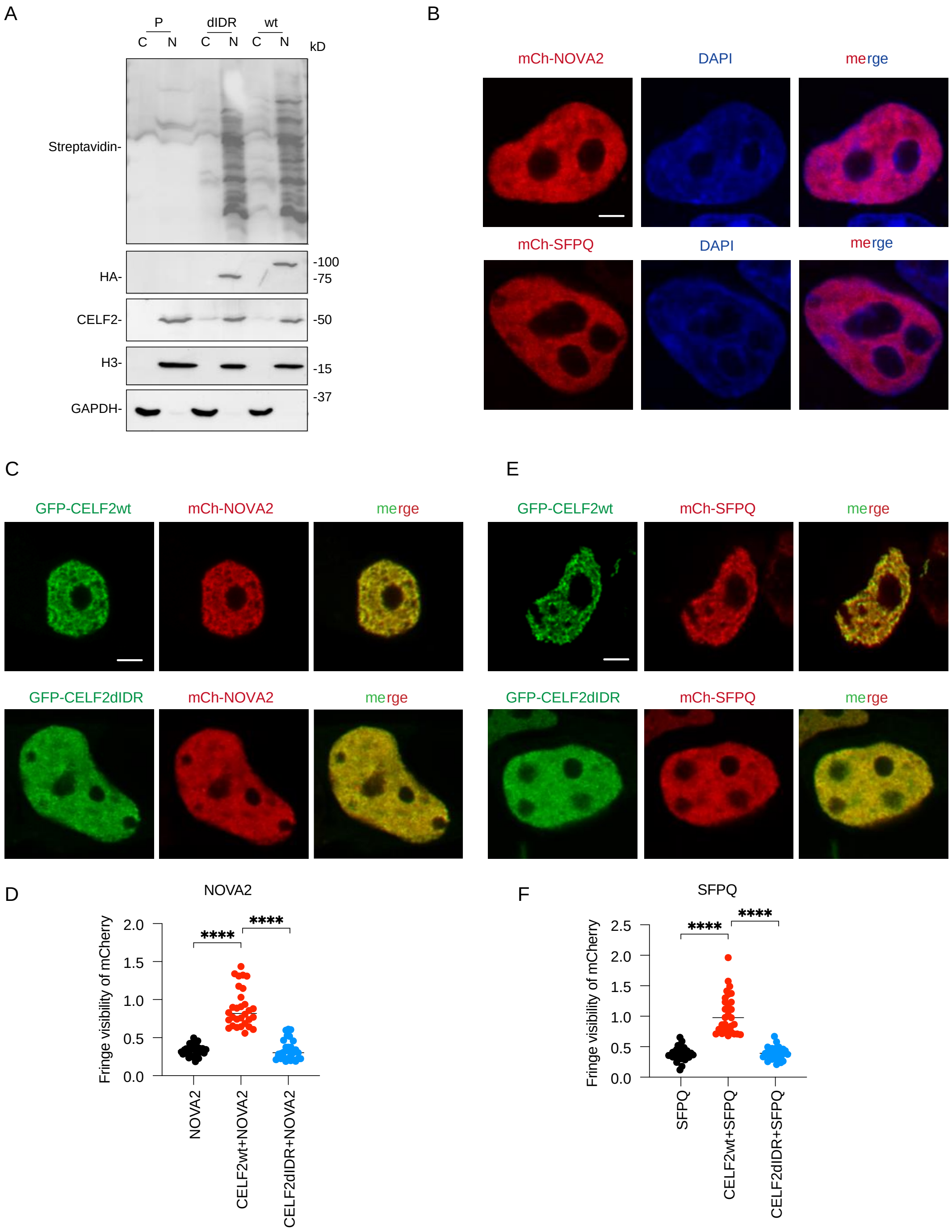


**Fig. S4. TurboID-identified cofactors of CELF2 colocalize with CELF2 in nuclear condensates and the colocalization is dependent on CELF2 IDR (related to Fig. 4).**
**(A)** Western blots confirming the inducible expression and *in vivo* proximity biotinylation in the established tet-on stable SH-SY5Y cell lines for CELF2 TurboID (wt and ΔIDR). The fractionation of nuclear fractions and loading amount were confirmed with Western blots for Histone H3

(nucleus-specific marker). The doxycycline-induced HA-BirA*-CELF2 fusion protein expression was detected by antibodies recognizing CELF2 and HA respectively. Various proteins biotinylated by HA-BirA*-CELF2 were detected using streptavidin-HRP blot.
**(B)** Representative images of mCherry-NOVA2 and mCherry-SFPQ in 293 cells, which have low endogenous CELF2 expression. mCherry-NOVA2 and mCherry-SFPQ displayed weakly punctuated expression pattern in the nucleus.
**(C-D)** Representative images of 293 cells co-transfected with mCherry-NOVA2 and GFP-CELF2 (wt, dIDR) and quantification of fringe visibility of mCherry+ condensates. When co-expressed with GFP-CELF2wt (but not GFP-CELF2ΔIDR), mCherry-NOVA2 exhibited a more punctate patten than when expressed alone. This suggests that NOVA2 might be recruited to CELF2 condensates through CELF2 IDR.
**(E-F)** Representative images of 293 cells co-transfected with mCherry-SFPQ and GFP-CELF2 (wt, dIDR) and quantification of fringe visibility of mCherry+ condensates. When co-expressed with GFP-CELF2wt (but not GFP-CELF2ΔIDR), mCherry-SFPQ exhibited a more punctate patten than when expressed alone.
Scale bar: 5 μm. Statistics: one-way ANOVA, ****P < 0.0001.

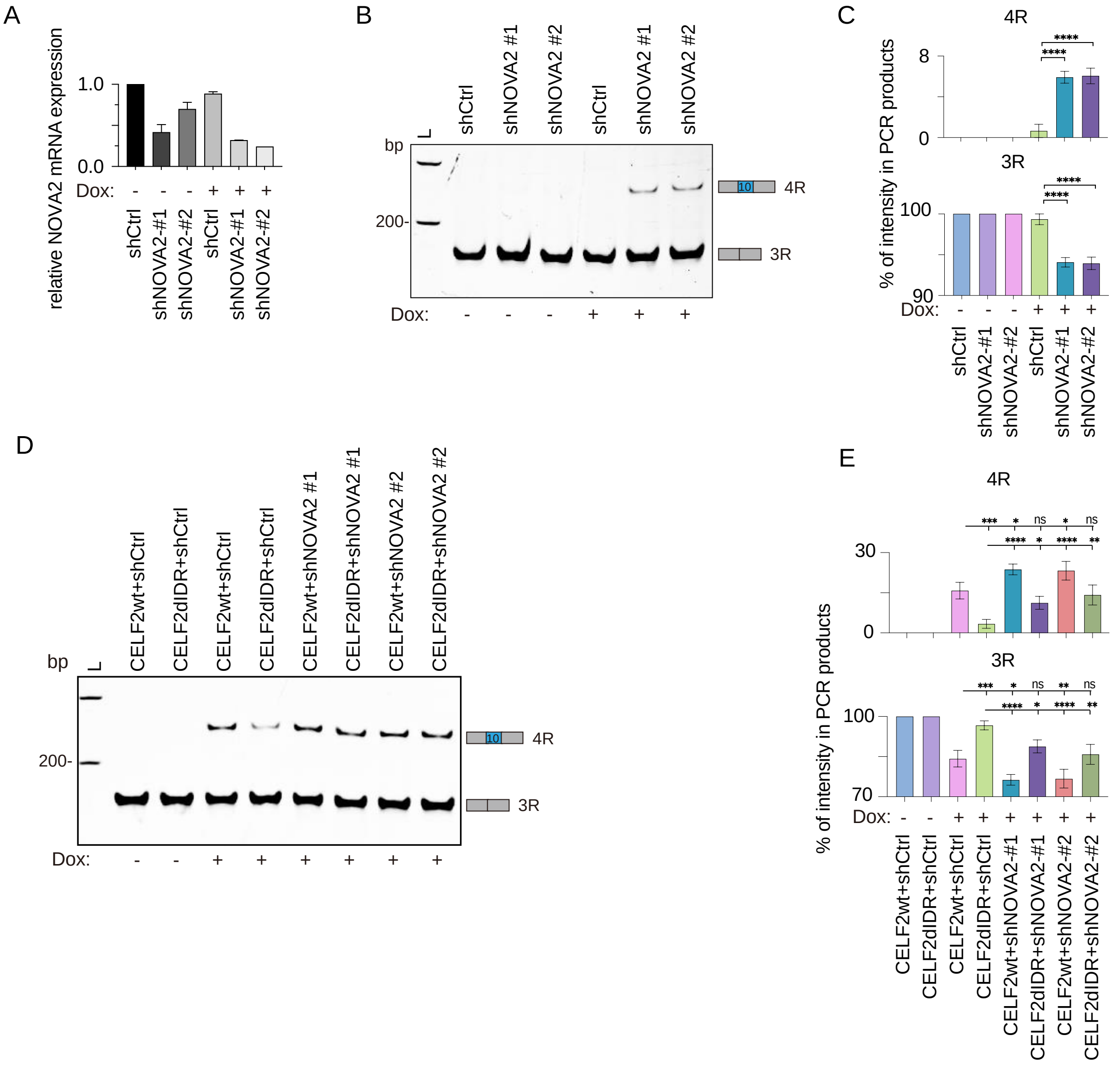


**Fig. S5. CELF2 and NOVA2 have opposing roles in regulating tau exon 10 splicing (related to Fig. 5).**

(**A**) RT-qPCR showing the knockdown efficiency of shNOVA2. Established stable SH-SY5Y cell lines with doxycycline-inducible expression of shNOVA2 were used to assess the efficiency of shRNA targeting NOVA2. Cells were treated with 100 ng/mL Doxycycline for 72 hours, followed by RNA extraction and RT-qPCR.

**(B-C)** Representative gel image and quantification of 4R and 3R tau transcripts in SH-SY5Y cells with NOVA2 knockdown (KD). Established stable SH-SY5Y cell lines with doxycycline-inducible expression of shNOVA2 were used to assess the alternative splicing of *MAPT* exon 10. Cells were treated with 100 ng/mL Doxycycline for 72 hours, followed by RNA extraction and RT-PCR. NOVA2 KD led to enhanced 4R tau expression, mirroring the effect of CELF2 overexpression.

**(D-E)** Representative gel image and quantification of 4R and 3R tau transcripts in SH-SY5Y cells with CELF2 overexpression and/or NOVA2 KD. Established stable SH-SY5Y cell lines with doxycycline-inducible expression of CELF2 and/or shNOVA2 were used to assess the alternative splicing of *MAPT* exon 10. Simultaneously overexpressing CELF2 and knocking down NOVA2 resulted in further enhanced 4R tau expression compared to single manipulation alone.

Statistical analysis: one-way ANOVA, *P < 0.05, **P < 0.01, ***P < 0.001, and ****P < 0.0001.

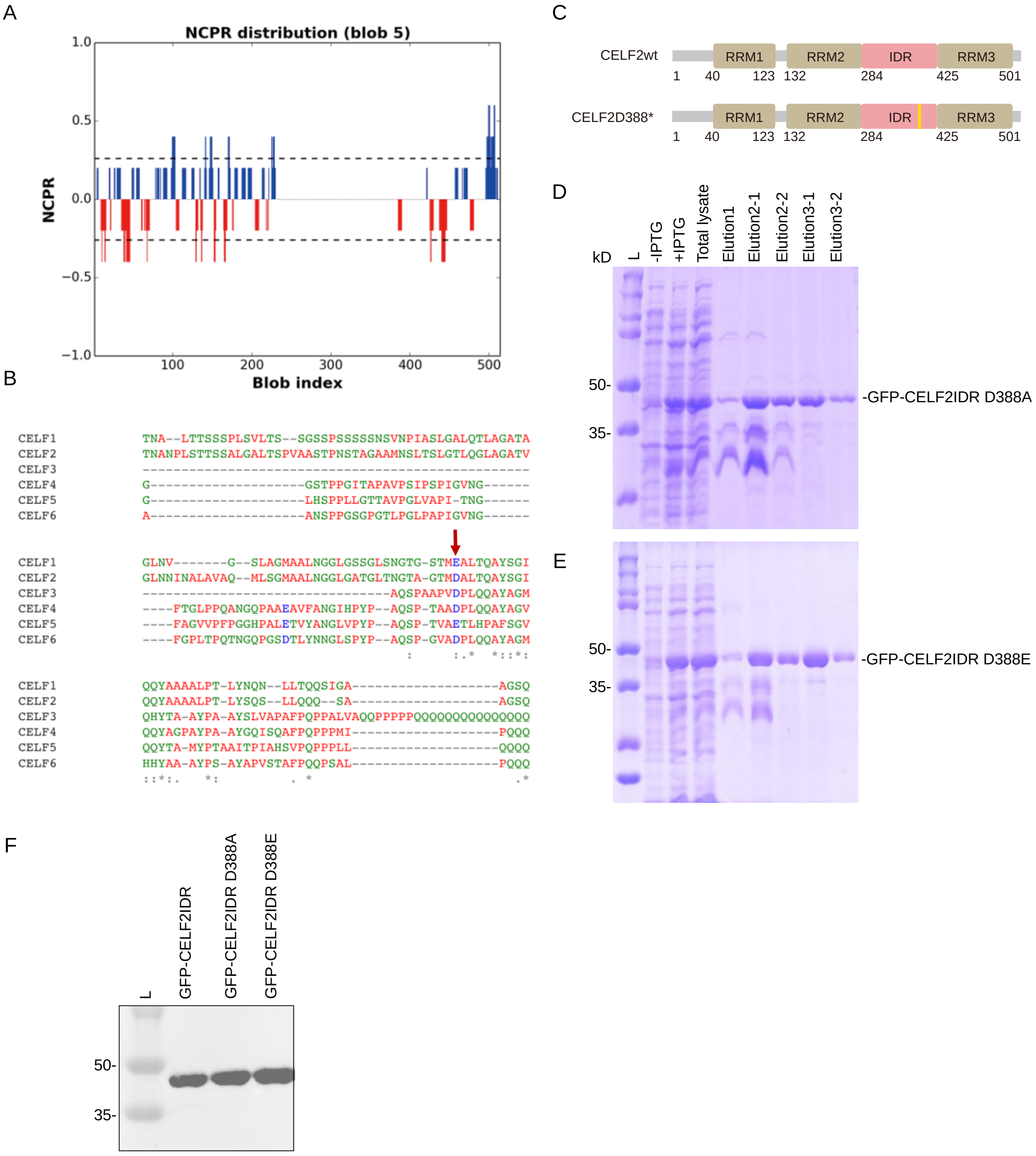


**Fig. S6. A conserved negatively charged residue within CELF2IDR is critical for CELF2 condensation and function (related to Fig. 6 and Fig. 7).**

**(A)** Net charge per residue (NCPR) for CELF2. There is only one charged residue (D388) within the entire CELF2IDR (hinge domain).

**(B)** Protein alignment of CELF hinge domains of the six CELF family members showing that the negatively charged residue (either D or E) within the hinge domain is conserved in all six CELF proteins.

**(C)** Schematic illustration of CELF2wt and CELF2D388 mutation.
**(D-E)** Coomassie staining of GFP-CELF2IDRD388A and GFP-CELF2IDRD388E from indicated protein samples. Prior to protein purification, we compared cell lysates from cells without and with IPTG induction. Total lysate from protein purification experiment, flow through and elutes from Ni+ columns were also examined.
**(F)** Western blot using anti-CELF2 antibody (recognizing the C-terminus of hinge domain) to confirm that the purified proteins are correct.
